# Recognition of A Highly Conserved DSRCPTQ Epitope in Envelope Protein of Zika Virus Through *in silico* Approaches

**DOI:** 10.1101/2020.02.11.943530

**Authors:** Anisur Rahman, Md. Iqbal Hossain, Saheda Tamanna, Md Neamat Ullah

## Abstract

The recurrent and recent outbreak of Zika Virus (ZIKV) has turned into global concern as yet there is no appropriate preventive measure been found. Situation getting worse as this virus also associates with several birth defects in neonatal such as primary microcephaly as well as many other neurological disorders. ZIKV adopts a wide host range which has hastened its expansion more recklessly. Hence, now there is an acute demand for developing a preventive vaccine against ZIKV. Immunoinformatic techniques have been employed in this study to pick out a highly conserved versatile antigenic B-cell linear epitope for all strains of ZIKV. Capsid protein (C), Membrane Glycoprotein Precursor (PrM), envelope protein (E) and RNA-dependent RNA Polymerase (NS5) have investigated by the implementation of sequence analysis and different epitope prediction methods. Some potential linear peptides have been recognized and tested for hydrophilicity and conservancy. Peptide with best antigenic properties was selected as ultimate final epitope and further structural exploration revealed its compatible position in protein 3D structure. Being fully conserved in all strains of ZIKV and its position in Envelope protein suggested epitope DSRCPTQ can be a quantum leap in the advancement of ZIKV prevention. However, extensive *in vitro* plus *in vivo* experimentations are needed to be clarified about the real potency of the selected epitope.

## 1 Introduction

Zika virus is a member of Flaviviridae virus family which contains a single-stranded positive-sense RNA (Malone et al., 2016). Its genome size is about 10.7 kb that encodes a single polyprotein that cleaved into different structural and non-structural proteins. Non-structural proteins (NS1, NS2A, NS2B, NS3, NS4A, NS4B and NS5) are associated with virus replication and multiplication in the host cell. And, structural proteins (capsid protein (C), Membrane Glycoprotein Precursor (PrM)-that cleaved into mature membrane protein (M) and envelope protein (E)) are involved in viral assembly (Lindenbach and Rice, 2003). The name of Zika Virus is based on Zika Forest of Uganda, where the virus was isolated from a rhesus macaque monkey for the first time in 1947 (Sikka et al., 2016).

First ZIKV infected human was noticed from a serological survey in Uganda (Dick, 1952). Later on, several studies in African and Asian countries revealed that the virus had been widespread within the human population of these regions (Wikan and Smith, 2016). In 2015 there as an outbreak of ZIKV in Brazil since then it has spread through tropical Americas and in 2015 and 2016 it became epidemic. The spread of ZIKV to other countries around the world is also reported in several studies (Duffy et al., 2009; Ety, 2014).

Though preliminarily it was believed that ZIKV causes mild feverish infection with rash and headache (Dick et al., 1952), now it has also been reported to associate with Guillian-Barr Syndrome (GBS) (Cao-Lormeau et al., 2016) and several other neurological complications as well (Tappe et al., 2015). Primary microcephaly and miscarriage in the time of pregnancy also have been noted in many cases due to ZIKV infection (Brasil et al., 2016; Mlakar et al., 2016). Besides, the presence of ZIKV in infected patients semen indicates that it might transmit by sexual means (Mansuy et al., 2016). Till date, no therapeutics or vaccine has been approved against ZIKV which makes its expansion and prevention more difficult (Abbink et al., 2018; Fernandez and Diamond, 2017; NIH, 2018).

In the recent years research on ZIKV have remarkably increased (Aid et al., 2017; Faye et al., 2014). Structure of the virion has been resolved and complete genome sequence was identified (Cunha et al., 2016; Kostyuchenko et al., 2016). ZIKV mainly targets neuroprogenitor cells, which eventually causes Congenital Zika Syndrome (CZS) and apoptosis (Lin et al., 2017). ZIKV is closely related with many other member of Flavivirus family such as dengue virus (DENV), West Nile virus (WNV), tick-borne encephalitis virus (TBEV) Japanese encephalitis virus (JEV) which advanced its vaccine development exclusively (Lanciotti et al., 2008).

Currently, a number of vaccines (for example; DNA vaccine, mRNA vaccine, purified inactivated virus vaccine, and viral vector vaccines) for ZIKV are in preclinical and clinical development. Plasmids-coding the gene of interest toinduce either of humoral and cellular immunity-are called DNA vaccine (Ferraro et al., 2011). Several ZIKV membrane glycoprotein precursor (PrM) and envelope protein coding DNA vaccines are still under clinical trial (Gaudinski et al., 2018; Tebas et al., 2017). About three ZIKV inactivated virus vaccine (ZPIV) were reported to be in phase I clinical trials (Modjarrad et al., 2018). mRNA vaccines are comparatively newer type of vaccine, which also been developed for ZIKV (Pardi et al., 2018). Additionally, viral vector-based approaches have been implemented to develop ZIKV vaccine. Various viruses (modified vaccinia virus Ankara (MVA), measles virus (MV) and adenovirus (Ad) vectors) are being engineered to function as ZIKV vaccine (Brault et al., 2017; Xu et al., 2018).

Researchers have been highly interested in Envelope protein (E), NS3 and NS5 as these proteins have a significant role in viral access and replication into the host cell (Malet et al., 2008; Perera et al., 2008). Studies on flavivirus vaccine revealed that antibodies designed to bind with Envelope protein (E) might be an efficient protection against ZIKV (Nowakowski et al., 2016). But, the risk of autoimmune response against potential epitope must be considered during vaccine development for ZIKV (Homan et al., 2016).

Various computer aided immunological approaches have been promising in vaccine design because of the availability of a large number of genomic and immunoinformatic data (De Gregorio and Rappuoli, 2012; Patronov and Doytchinova, 2013). Nowadays vaccine design has become more facile than ever in the matter of both time and expenses due to the application of computational methods and algorithms in this field.

Generally, Epitopes are two types; continuous and discontinuous. Continuous epitopes are simple linear peptide sequences while discontinuous epitopes are structurally more complex, nonlinear and assembled together for folding in the natural form of protein. According to their corresponding receptors, epitopes can be also classified into B- and T cell epitopes. B cell epitopes contain both continuous (10%) and discontinuous (90%) epitopes, while most of T cell epitopes are continuous. Antibodies or B-cell receptors recognize B-cell epitopes. Four approaches have been employed for epitope prediction: sequence-based methods, structure-based methods, hybrid methods, and consensus methods. Though the hybrid method provides more accurate results than others, consensus methods give the best prediction results (Yang and Yu, 2009).

Though some investigations appeared to be auspicious in preclinical and clinical trials, no vaccine have been approved for use. The objectives of this computational approach were to (1) investigate the recent advancement in vaccine development against ZIKV thoroughout the world, (2) identify highly conserved region in ZIKV genome as this virus confer high versatility amongst different strains and region.

The main aim of the study was to suggest at least one precise epitope for ZIKV vaccine development, which would be universal for all strains of this virus with immunogenic properties.

In current investigation, a B-cell epitope *DSRCPTQ* has been discovered from the sequence of ZIKV which is 100% conserved in all strains from all region and hosts of this virus. Several sequence analyses, such as Multiple sequence alignment (MSA), retrieval of the most consensus sequence for ZIKV, determination of the conserved region as well as structural validation such as epitope mapping on protein 3D structure were accomplished to predict highly immunogenic, surface accessible and conserved epitope. Various computer programs and online server were implemented for our study.

## 2 Materials and Methods

### 2.1 Retrieval of Protein Sequences

Primary amino acid sequences of the complete genome for full polyprotein of Zika Virus were collected using Virus Variation tool (https://www.ncbi.nlm.nih.gov/genome/viruses/variation) of NCBI (Hatcher et al., 2017). Only full-length sequences have been selected and fully identical sequences were excluded. All other filter options (Host, Region/Country, Genome Region, Isolation Source) were set to default. Thus, a sum of 396 protein sequences has been collected which includes all available strains of Zika Virus in Virus Variation database from all over the world. Sequences with ambiguous or exceptional amino acid code (e.g., B, J, O, U, Z, X, –) causes error during epitope conservancy analysis. Thereupon, sequences containing such codes were excluded from the list. Finally, total 305 full-length protein sequences were chosen for further analysis.

### 2.2 Identification of Conserved Region

UGENE v1.31.1 (http://ugene.net) software package was employed for Multiple Sequence Alignment (MSA) (Okonechnikov et al., 2012). In this software, MSA was done by using ClustalO program (Sievers et al., 2011). From this Multiple sequence alignment, the most consensus sequence of full-length polyprotein of ZIKV was retrieved and protein sequence region of Capsid Protein (C), Membrane Glycoprotein Precursor (PrM), Membrane Glycoprotein (M), Envelope Protein (E), RNA-dependent RNA Polymerase (NS5) were checked for conserved segments (Table-1). Sequences with the highest number of identical and similar amino acid and no gap were selected as conserved regions. Total 12 conserved regions were picked to use for epitope prediction (Table-2).

**Table 1:** Positions of Proteins in Consensus Sequence from Multiple Sequence Alignment (MSA)

| From the Region of Protein | Position in Consensus Sequence from MSA |
| --- | --- |
| Capsid Protein (C) | 3-106 |
| Membrane Glycoprotein Precursor (PrM) | 125-292 |
| Membrane Glycoprotein (M) | 218-292 |
| Envelope Protein (E) | 293-792 |
| RNA-dependent RNA Polymerase (NS5) | 2523-3425 |

**Table 2:**
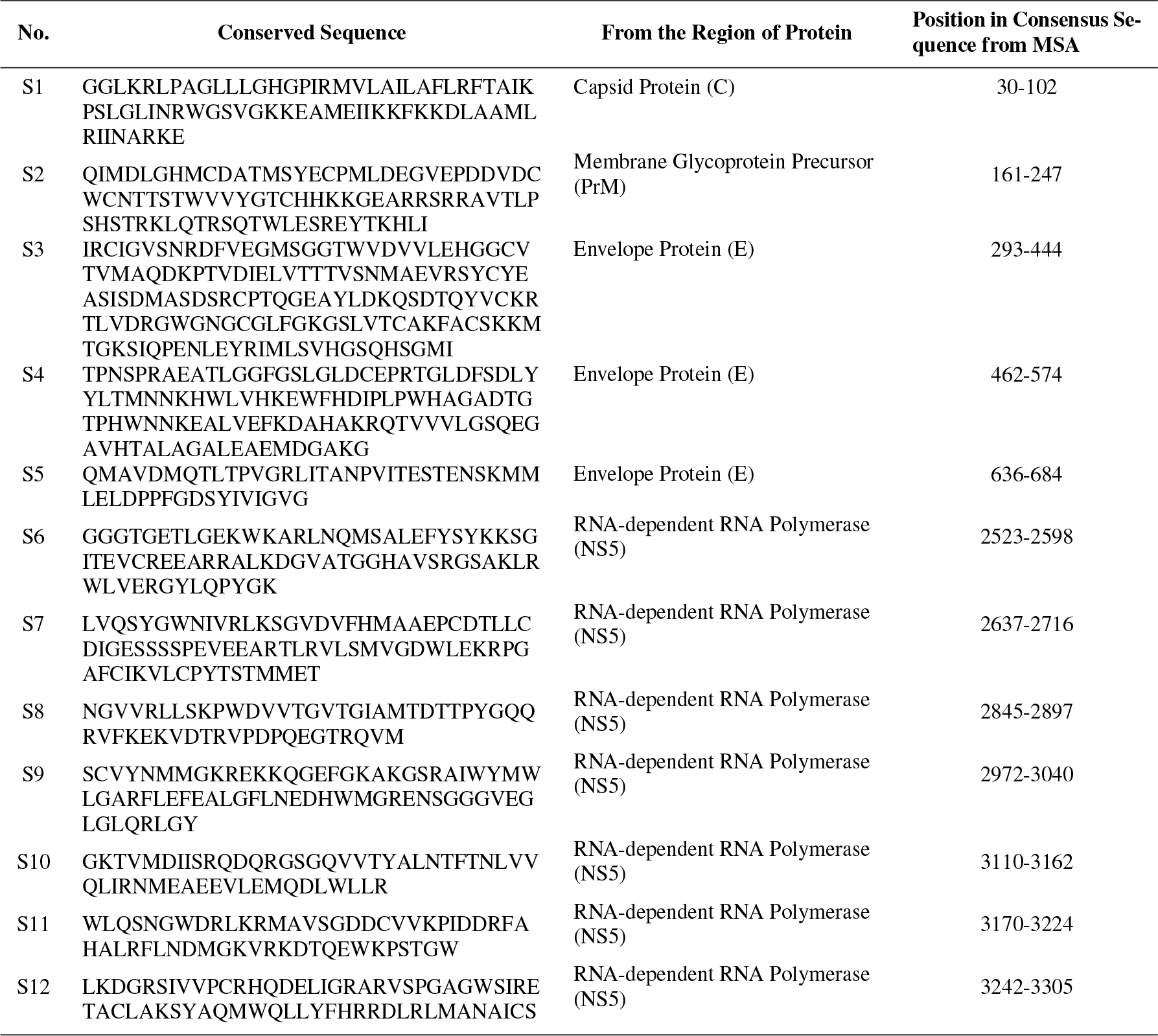
Conserved Sequence from consensus protein identified by Multiple Sequence Alignment (MSA) in UGENE software

### 2.3 Epitope Prediction

B-cell epitope prediction tool of the Immune Epitope Database (IEDB) was implemented to predict linear epitopes from selected conserved protein sequences (http://tools.iedb.org/main/bcell). Initially, the following three methods from this tool have been applied for linear epitope prediction.

1. *Bepipred Linear Epitope Prediction 2.0:* This method predicts B-cell epitopes from a protein sequence, using a Random Forest algorithm trained on epitopes and non-epitope amino acids determined from crystal structures (Jespersen et al., 2017). This method was used with the default threshold value of 0.5.
2. *Kolaskar & Tongaonkar Antigenicity:* For predicting epitopes based on antigenicity this method was employed with a threshold value of 1.0 (Kolaskar and Tongaonkar, 1990). Kolaskar & Tongaonkar Antigenicity scheme uses physicochemical properties of amino acid residues and their frequencies of occurrence in protein sequences to predict B-cell epitopes. The method can predict antigenic determinants with about 75% accuracy which is ever best-known method.
3. *Emini Surface Accessibility Prediction:* Then, Emini Surface Accessibility Prediction method from IEDB tool was utilized for predicting epitope based on surface accessibility scale with the threshold value of 0.8 (Emini et al., 1985).

A number of peptides from each conserved region were generated for each method. Only peptides with overlapping between three prediction methods were picked for further analysis.

#### 2.3.1 Prediction of Epitope Conservancy

To calculate the Conservancy of selected epitopes, the IEDB epitope conservancy analysis tool (http://tools.iedb.org/conservancy) was applied (Bui et al., 2007). This tool measures the degree of conservancy of an epitope within given protein sequences based on sequence identity.

#### 2.3.2 Prediction of Hydrophilicity

Parker Hydrophilicity Prediction method was employed to determine Hydrophilicity of conserved regions containing candidate epitopes (Parker et al., 1986). This method uses the average score as default threshold value for calculation. In this study default threshold value for every conserved sequence were kept unchanged.

### 2.4 Evaluation in Protein 3D Model

Phyre2 server (Protein Homology/analogy Recognition Engine V 2.0) was used to identify the 3D structure of protein E (Kelley et al., 2015). Based on homology modeling by Phyre2 (http://www.sbg.bio.ic.ac.uk/phyre2/html) chain A of protein 5IRE was found in PDB Database (https://www.rcsb.org) with 100% confidence and coverage (Sirohi et al., 2016). Ramachandran plot for modeled protein was done by employing Uppsala Ramachandran Server (http://eds.bmc.uu.se/ramachan.html) to analyze its the validity and quality (Kleywegt and Jones, 1996). Later on, the protein structure has been processed, and the conserved epitope was mapped on protein 3D structure by the use of Discovery Studio 4.5 Visualizer software (http://www.accelrys.com) (Dassault Systèmes BIOVIA, 2017).

## 3 Results

### 3.1 Selection of Conserved Regions

Multiple sequence alignment of full-length polyprotein of Zika virus have been checked for conserved peptide segments in the region of Capsid Protein (C), Membrane Glycoprotein Precursor/Membrane Glycoprotein (PrM/M), Envelope Protein (E), RNA-dependent RNA Polymerase (NS5) (See, Supplementary Materials: Sp1 to Sp4). And thus, 1, 1, 3 and 7 conserved peptide segments were identified in the region of C, PrM, E, NS5 protein respectively (Table-1). Total 12 conserved regions have been considered for epitope analysis. In UGENE a most consensus protein sequence of Zika virus polyprotein was found after performing MSA and conserved sequences were taken from this consensus sequence.

### 3.2 Prediction and Selection of B-cell Epitopes

To be an effective epitope, a peptide must confer immunogenic, antigenic and surface accessible properties. A competent B-cell epitope must have the surface accessibility property to bind with an antibody effectively (Caoili, 2010). Three methods (Bepipred Linear Epitope Prediction 2.0, Kolaskar & Tongaonkar Antigenicity and Emini Surface Accessibility Prediction) from the IEDB database have been utilized to recommend linear B-cell epitope from all conserved regions.

For every conserved amino acid sequence, a list of predicted peptides was generated in each method. Peptides which were common in all three methods were chosen as candidate epitope for further analysis. No epitope was found from conserved sequences in the region of Capsid Protein (Conserved Sequence-S1) and Membrane Glycoprotein Precursor (Conserved Sequence-S2). In envelope protein region, one epitope was identified from Conserved Sequence 3 (S3). Another, two epitopes were found from Conserved Sequence 6 and 12 (S6 and S12) in the NS5 protein region. All predicted results are listed in supplementary material Sp5.

1. *Envelope Protein (E):* Epitopes predicted by Bepipred 2.0, Antigenicity, and Surface Accessibility methods from Conserved Sequence 3 (S3) of Envelope Protein (E) region are listed in Table-3. An antigenic epitope *DSRCPTQ* was overlapped with Bepipred predicted peptide and surface accessible peptide *MASDSRCPTQGEAYLDKQSDT*. As this peptide fulfilled all three preliminary criteria to be a potential epitope, it was considered as a candidate epitope.
2. *RNA-dependent RNA Polymerase (NS5):* In this protein region 7 conserved amino acid segments (S6 to S12) were extracted. Every segment was analyzed for B-cell epitope prediction and two potential epitopes have been found out in S6 and S12. All predicted peptides from conserved region 6 from each method (Bepipred 2.0, Antigenicity, and Surface Accessibility) are summarized in Table-4. In this case, a peptide *FYSYKK* was found to be overlapped among other prediction results. So, this peptide was considered as a candidate B-cell epitope.

**Table 3:**
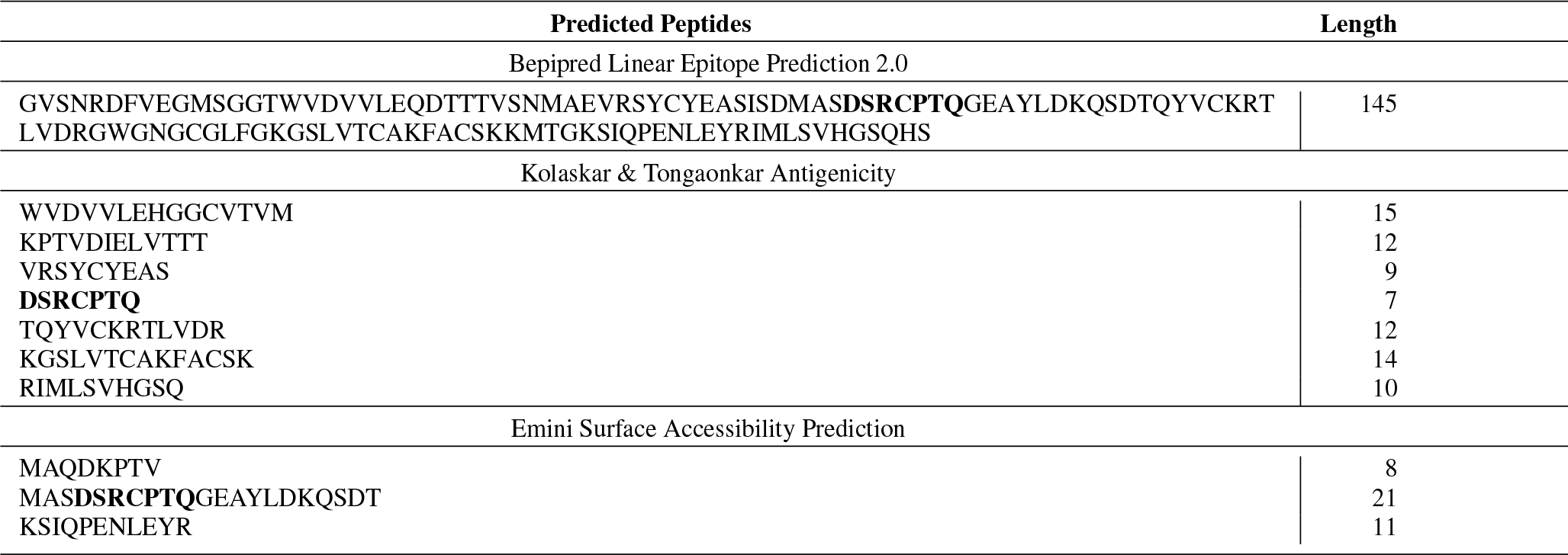
Bepipred Linear Epitope, Antigenic and Surface Accessible epitopes predicted from Conserved Region S3 of Envelope Protein (E)

**Table 4:**
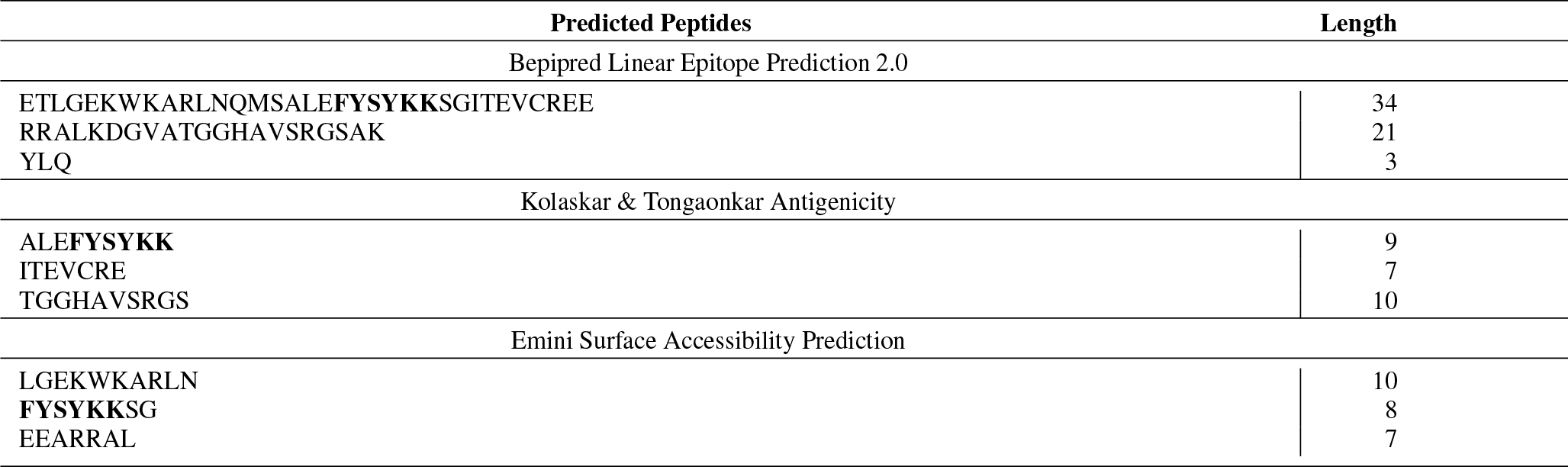
Bepipred Linear Epitope, Antigenic and Surface Accessible epitopes predicted from Conserved Region S6 in RNA-dependent RNA Polymerase (NS5)

Lastly, another candidate epitope *RHQDE* has been identified from S12 region on NS5 protein. This peptide also found to be overlapped with all three predictions. All predicted epitopes from Conserved Sequence 12 are listed in Table-5.

**Table 5:** Bepipred Linear Epitope, Antigenic and Surface Accessible epitopes predicted from Conserved Region S12 in RNA-dependent RNA Polymerase (NS5)

| Predicted Peptides | Length |
| --- | --- |
| Bepipred Linear Epitope Prediction 2.0 |  |
| RSIVVPC**RHQDE**LIGRARVSPGAGWSIRETAC**LAKSY**AQMW | 41 |
| DLRLMAN | 7 |
| Kolaskar & Tongaonkar Antigenicity |  |
| RSIVVPC**RHQDE** | 12 |
| RARVSPG | 7 |
| ETAC**LAKSY** | 9 |
| MWQ**LLYFHR** | 9 |
| Emini Surface Accessibility Prediction |  |
| **RHQDE**L | 6 |
| FHRRDL | 6 |

#### 3.2.1 Conservancy Analysis

Epitope Conservancy Analysis tool of IEDB has been implemented to find out the percentage of conservancy of all candidate epitope *(DSRCPTQ, FYSYKK and RHQDE)*. All epitopes demonstrated 100% conservancy amid all 305 sequences of Zika virus (Figure-1).

**Fig. 1:**
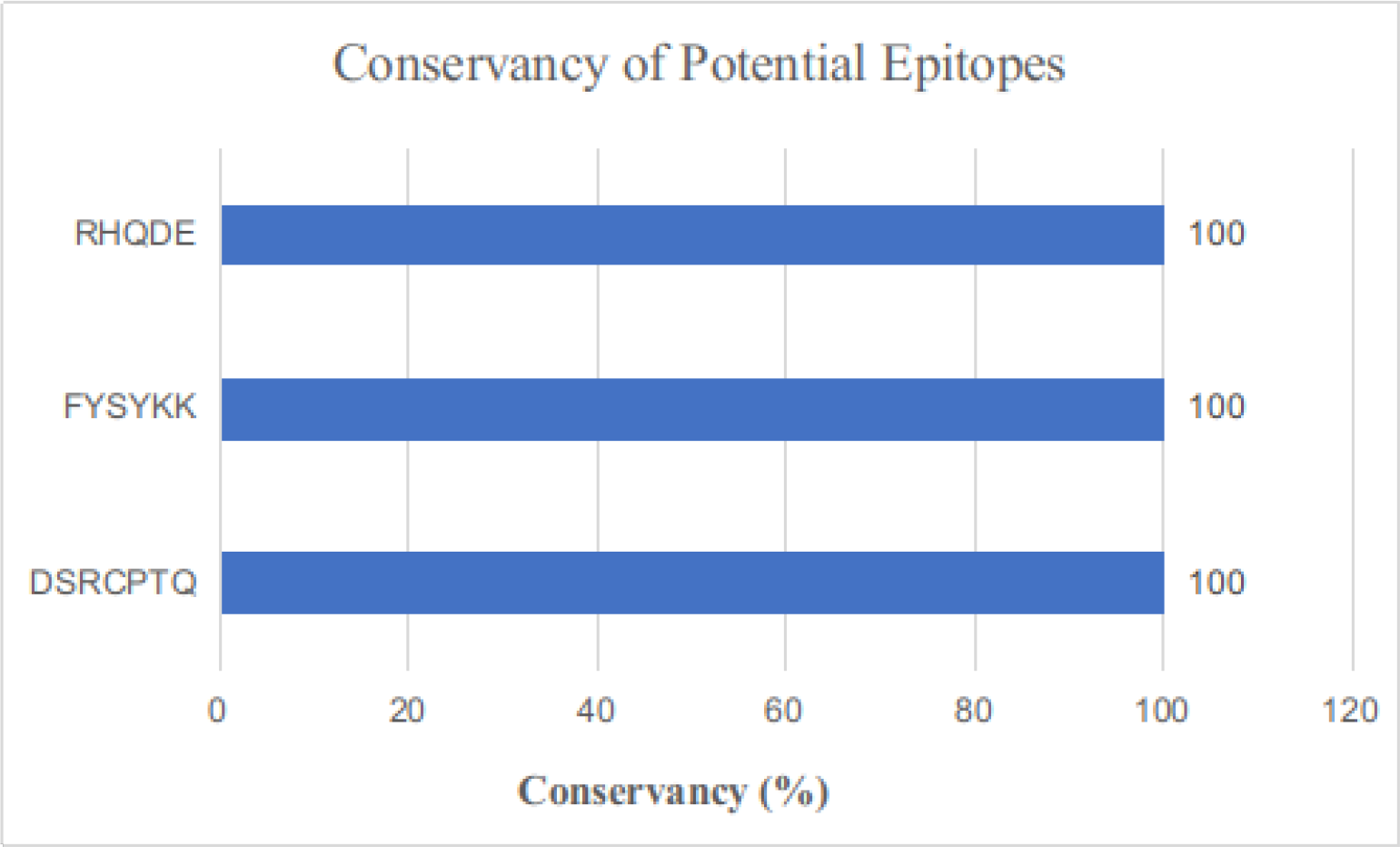
Results of Conservancy Analysis for three predicted epitopes

#### 3.2.2 Hydrophilicity of Epitopes

Potential epitopes containing conserve sequence S3, S6, S12 were tested for hydrophilicity property using Parker Hydrophilicity Prediction method. Hydrophilicity of these three conserved sequences is demonstrated in Figure-2. Average hydrophilicity score of each sequence was used as default threshold value. For S3 sequence default threshold value was 2.137 and residue score of predicted epitope *DSRCPTQ* was 5.057, that is much higher than default threshold value. Average hydrophilicity of S6 sequence was 2.005 but predicted epitope *FYSYKK* from this region had much lower residue score (0.817). Though predicted epitope *RHQDE* from sequence S12 showed higher residue score 6.02, threshold value of this conserved region was very low 0.738 and length of this epitope is also quite short to be a suitable epitope. Summary of Parker Hydrophilicity results for all three epitopes and corresponding conserved sequences are tabulated in Table-6.

**Fig. 2:**
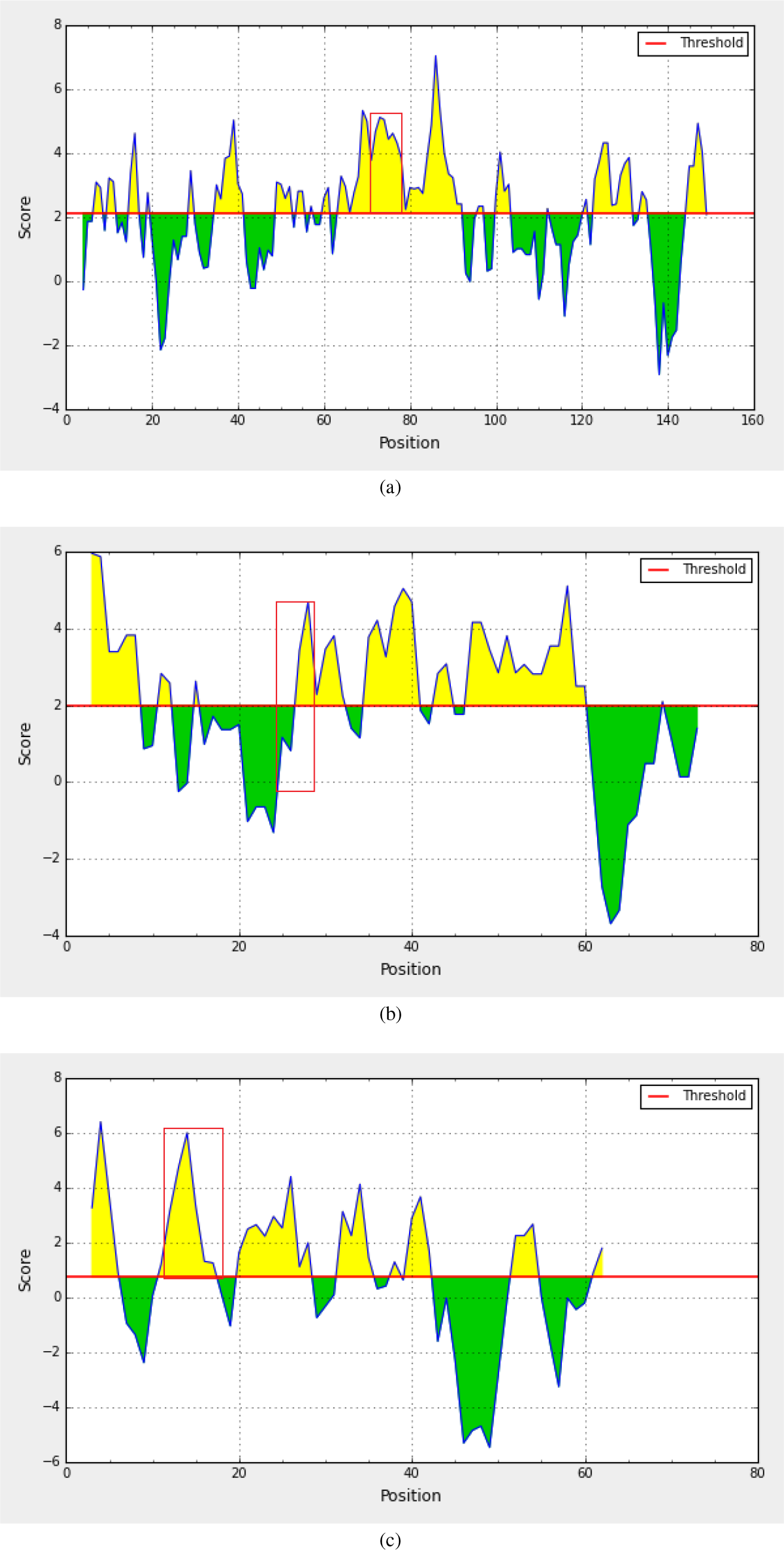
Hydrophilicity plot of (a) conserved sequence S3 from Envelope protein, (b) S6 and (c) S12 from NS5 protein. In every graph yellow color represents hydrophilic regions. Red boxes represent the hydrophilicity of candidate epitope from corresponding conserved sequence

**Table 6:** Parker Hydrophilicity results of candidate epitopes

| Epitope | Conserved Sequence | Protein | Threshold Value of Conserved Sequence (Average Score) | Residue Score |
| --- | --- | --- | --- | --- |
| DSRCPTQ | S3 | Envelope Protein (E) | 2.137 | 5.057 |
| FYSYKK | S6 | RNA-dependent RNA Polymerase (NS5) | 2.005 | 0.817 |
| RHQDE | S12 | RNA-dependent RNA Polymerase (NS5) | 0.738 | 6.02 |

Considering all properties and criteria, ***DSRCPTQ*** in Envelope protein sequence was chosen as final desired B-cell epitope.

### 3.3 Epitope Mapping on Protein 3D Structure

According to the result from Ramachandran plot analysis 84.3% amino acid residues of ZIKV Envelope protein are in the acceptable region (Figure-3). The epitope ***DSRCPTQ*** was found to be located on the surface of the 3D structure of Envelope Protein after mapping (Figure-4).

**Fig. 3:**
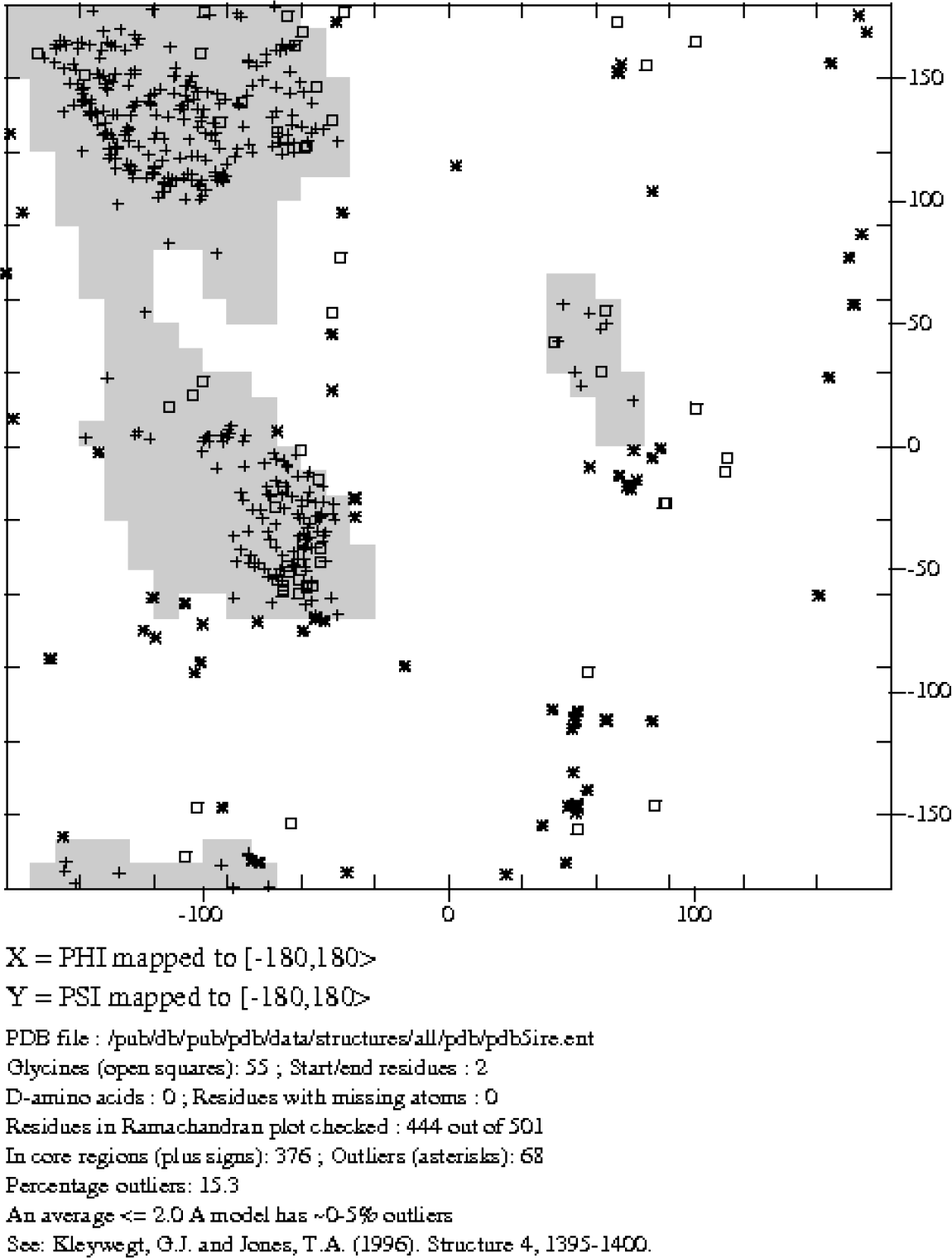
Ramachandran plot for Envelope Protein by Uppsala Ramachandran Server

**Fig. 4:**
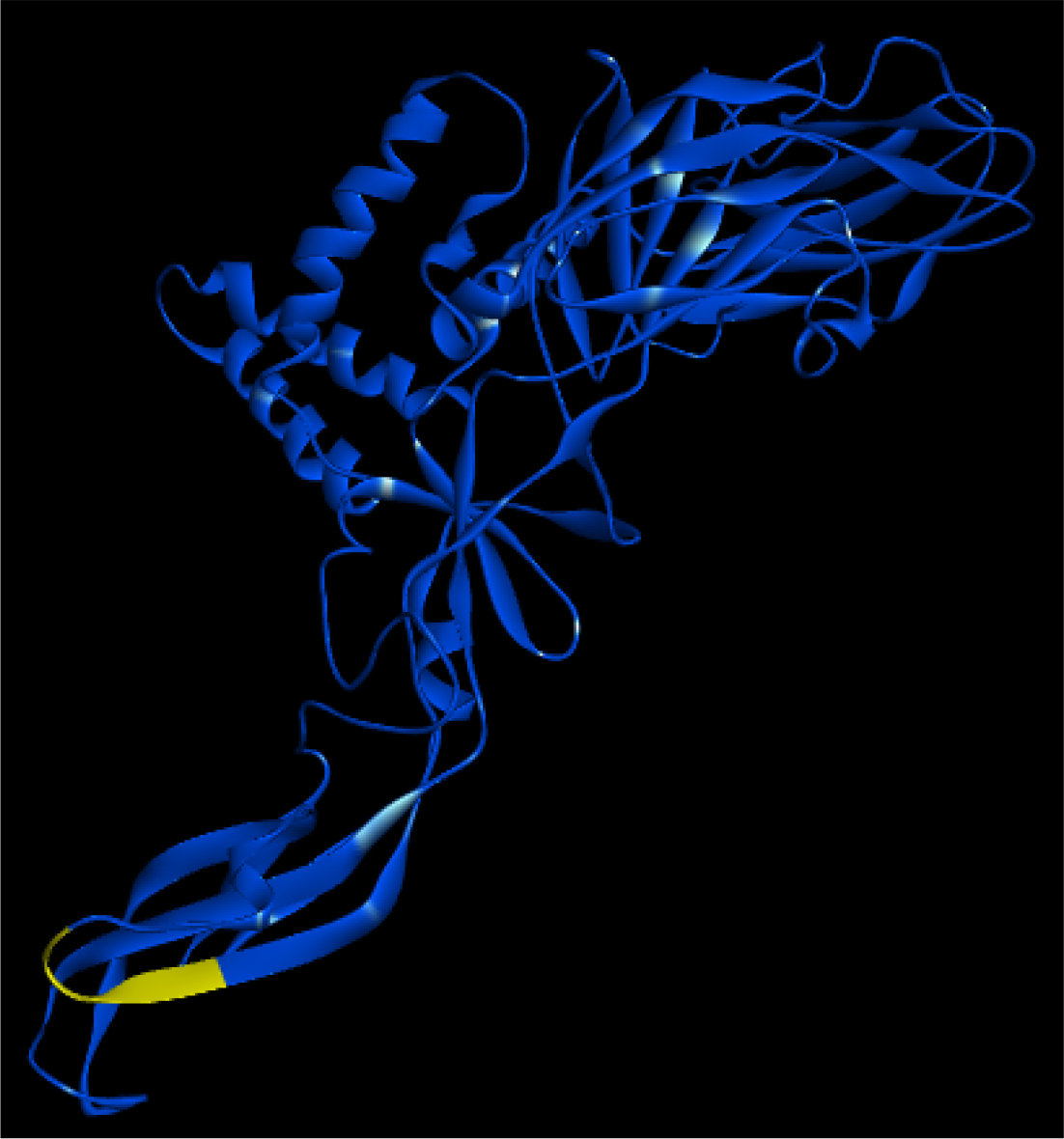
Epitope Mapping on E Protein 3D structure

## 4 Discussion

Zika virus epidemic has become a global threat in recent year. Over the past few decades, this virus has transmitted actively (local transmission) around different countries and territories (Countries, 2016). Zika virus uses a wide number of organisms as host including human, mosquito (mainly Aedes genus), monkeys and as well as other animals especially in several mammals (Olson et al., 1983; Wang et al., 2016). Thats why full polyprotein sequences of all strains of Zika virus around the world from all listed hosts in virus variation were included in this study.

According to the World Health Organization vaccine for Zika virus should be inactivated and non-live as pregnant women are highly susceptible to this virus (Rasmussen et al., 2016). So, the peptide-based vaccine would be best effective preventive measure against this virus. Protein E and PrM coding plasmid DNA vaccine has been developed which will encode the outer protein coat of Zika virus virion (Dowd et al., 2016). Flavivirus envelope glycoproteins play a crucial role in the initiation of endocytosis into the host cell by binding with endosomal membrane (Dai et al., 2016). So, an antibody against envelope protein of Zika virus will be effective to prevent the entrance of virus into the host cell. Until July 2018, there is no appropriate vaccine for Zika virus has been approved for clinical use, though some vaccines are currentlyin the clinical trial (NIH, 2018). So, it is a crying need to construct a suitable preventive vaccine against this global threat.

*in silico* methods have been turning into popular and essential tools in vaccine model as it helps to curtail the time and the number of in vitro laboratory assays (Backert and Kohlbacher, 2015; Florea et al., 2003; Groot and Rappuoli, 2004). Now scientists are following various immunoinformatic tools as cost-efficient and rapid approaches to develop the vaccine against Zika virus, for example, antigenic epitope prediction from Zika proteins (Alam et al., 2016; Ashfaq and Ahmed, 2016; Badawi et al., 2016; Dar et al., 2016; Khan et al., 2015; Weltman, 2016). In this research work, a very potential epitope in the sequence of envelope protein was found which is universal for all the strains of Zika virus all over the world.

From MSA of full-length polyprotein of Zika Virus, several conserved regions (S1-S12) were selected in the coding sequence of Capsid Protein (C), Membrane Glycoprotein Precursor (PrM), Envelope Protein (E), RNA-dependent RNA Polymerase (NS5). The most consensus sequence of full-length polyprotein among all Zika Virus strains also been found from MSA result in UGENE software and conserved segments were taken from this consensus sequence. After analysis, each conserved segment in terms of Bepipred linear epitope prediction, antigenicity and surface accessibility three epitopes from conserved segment S3, S6 and S12 were identified, one from each segment (Table-6). Though all epitopes were 100% conserved in all sequence of Zika virus, epitope *DSRCPTQ* from conserved region S3 possesses better hydrophilicity property based on Parker Hydrophilicity Prediction method (Table-6). Additionally, the presence of S3 conserved segment in envelope protein sequence makes the epitope more acceptable.

Furthermore, after structural analysis by epitope mapping on Envelope protein 3D structure the epitope was located on the external surface of the protein, thereby it added further affirmation that *DSRCPTQ* epitope is endowed with all the compulsory properties to be a suitable B-cell epitope. Principle of virus detection through molecular immunoassay techniques is based on conserved epitopes of particular viral protein. Based on this principle Zika virus Envelope protein is of great interest for epitope detection among several researchers (Weltman, 2016).

As only computational approaches were implemented to identify this epitope, additional *in vivo* and *in vitro* evaluation is also required to determine the real efficacy, antigenicity and other immunogenic properties of the epitope. An additional adjuvant can be coupled with this peptide to enhance its immunogenicity and stability (Gershoni et al., 2007).

Defining the antigenic sites in viral protein is the fundamental basis for the generation of therapeutics and vaccine against viral diseases (Toyoda et al., 2000). By adopting sequence analysis and various computational prediction method *DSRCPTQ* peptide was identified from Zika virus envelope protein (E) as the best epitope to design new specific common antibody for all strains of this virus upon *in vitro* and *in vivo* test.

## 5 Conclustion

As ZIKV spreading throughout the world very rapidly and deadly, it is an immediate call for discovery of a standard universal vaccine, which will be effective against all strains of this virus. A number of researchers have been working tirelessly to develop preventive therapeutics and vaccine against this fatal virus and several remarkable outcomes also been noticed. This study was an effort to take this endeavor one step forward. A potential epitope has been identified in the envelope protein region of ZIKV which is conserved in all strains. To avert the further outbreak of ZIKV, vaccine, and therapeutics based on this suggested epitope can be an efficient arsenal. As it was an *in silico* approach consequential *in vitro* and *in vivo* validations are also necessary.

