## Supplementary figures and images for "Recognition of A Highly Conserved DSRCPTQ Epitope in Envelope Protein of Zika Virus Through *in silico* Approaches"

### Supplementary Material 1

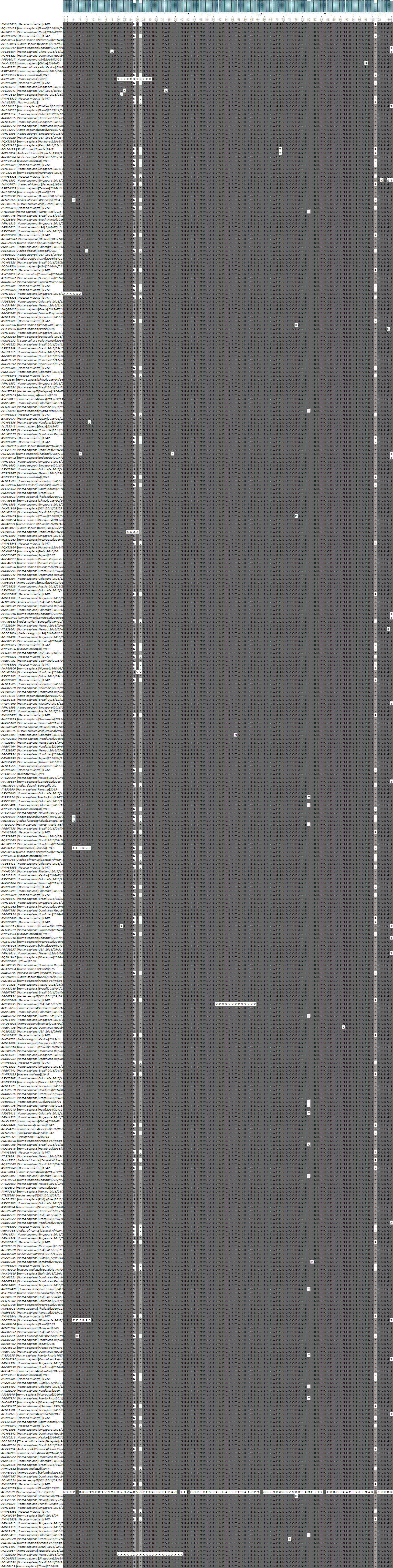

### Supplementary Material 2

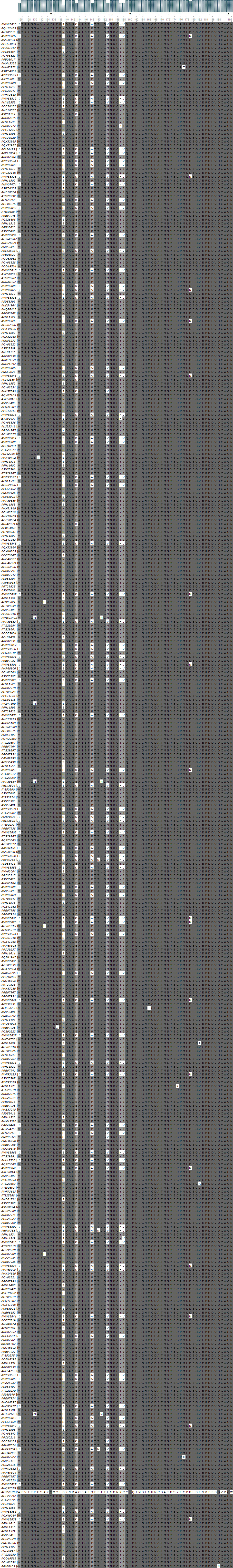

### Supplementary Material 3

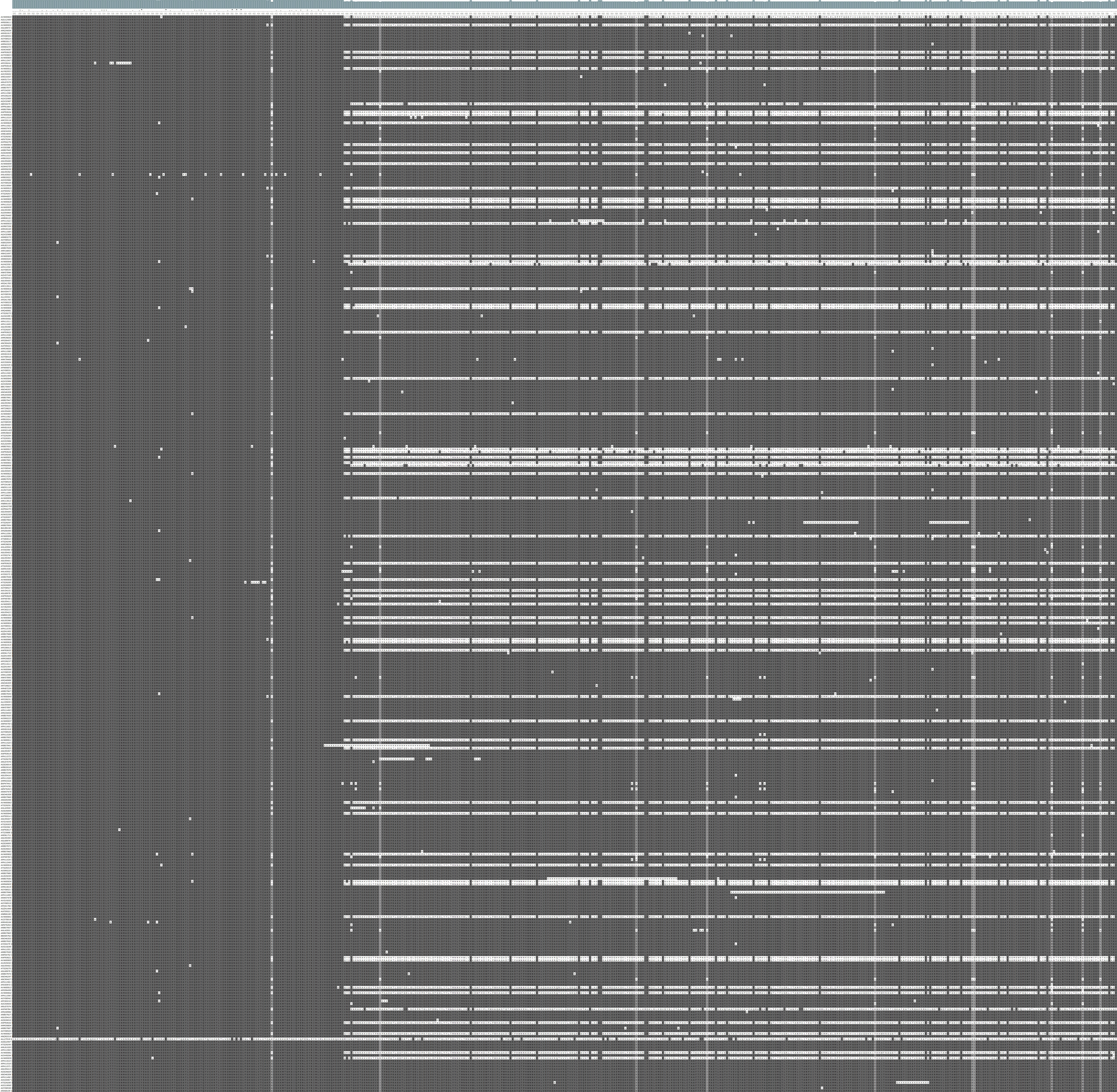

### Supplementary Material 4

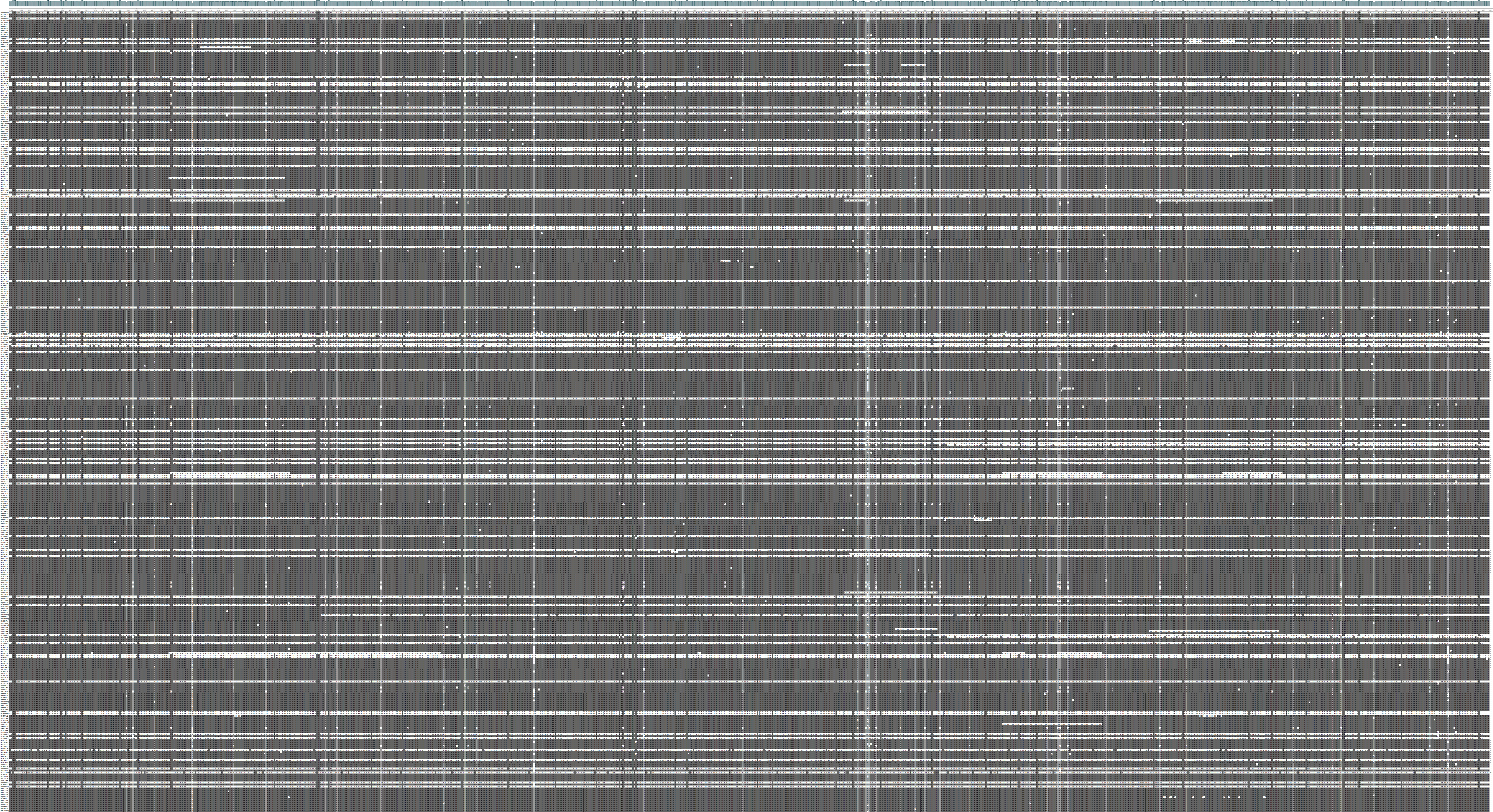
